# Molecular signatures of heat tolerance in an Australian alpine plant during moderate warming

**DOI:** 10.1101/2023.09.27.559694

**Authors:** Rocco F. Notarnicola, Jemimah Hamilton, Pieter A. Arnold, Ashley W. Jones, Stella Loke, Zhi-Ping Feng, Alexander N. Schmidt-Lebuhn, Benjamin Schwessinger, Adrienne B. Nicotra, Diep R. Ganguly

## Abstract

Understanding the molecular mechanisms of heat tolerance could help to predict the impacts of climate change on our native flora. However, much of our current understanding is derived from *Arabidopsis thaliana* and select crops exposed to short and intense periods of heat stress. Here, we characterise the transcriptomic response of *Wahlenbergia ceracea*, a perennial herb found in Australia’s subalpine regions, to sustained moderate warming. We contrasted responses between relatively heat-tolerant and heat-sensitive lines grown under cool (24/15 ºC day/night) or warm (30/20 ºC) temperatures. We observed that sustained warming up-regulated genes related to RNA regulation, including modification and splicing, and down-regulated genes associated with photosynthesis and plastid organization. Interestingly, heat-tolerant lines demonstrated a more pronounced repression of photosynthesis-associated genes, compared to heat-sensitive lines, suggesting that the regulation of the light-harvesting machinery may contribute to photosystem thermal tolerance. Co-expression analyses revealed only weak module-level correlations with direct measures of thermal tolerance. However, stronger associations were evident with chlorophyll content and photosynthetic efficiency, which might indirectly influence thermal tolerance. Prediction of transcription factors targeting warming-responsive genes implicated hormone signaling networks, especially ethylene, in contributing to differences in thermal tolerance. In conclusion, we present genomic resources for transcriptome analyses in *W. ceracea* and highlight the contrasting gene regulatory patterns between relatively heat-tolerant and heat-sensitive *W. ceracea* lines experiencing sustained moderate warming.

## Introduction

Alpine environments, which are characterised by high spatio-temporal variability in climatic conditions including steep temperature and precipitation gradients across complex topography, are predicted to be severely impacted by global warming (Gobiet et al., 2014). Australia’s alpine and subalpine regions harbour many plant species that are vulnerable to climate change (Sritharan et al., 2021). Plants occupying distinct elevations exhibit differing levels of phenotypic plasticity, suggesting that there are distinct molecular processes regulating growth and reproduction (Nicotra et al., 2015). However, genomic and transcriptomic studies on native plant species occupying such environments remain under-represented (Geange et al., 2021). Such studies can uncover the genomic architecture and molecular basis for trait selection, their variation, and plasticity, including under changing environmental conditions (Oomen and Hutchings, 2017; Rashid et al., 2020; Oomen and Hutchings, 2022). These outcomes have the potential to inform conservation planning by, for example, determining population structure, defining conservation units, and constructing genomic reaction norms (Vandersteen Tymchuk et al., 2010; Oomen and Hutchings, 2022). Therefore, expanding genomic investigations into wild species, especially those that naturally occupy vulnerable environments, can produce valuable insights.

Understanding plant responses to heat is intensely pursued due to the projected increases in mean surface temperatures (+3 °C by 2100) as a result of global warming (IPCC, 2023). Many studies focus on short, intense heat events, referred to as heat-shock (Richter et al., 2010). Heat shock causes several changes to the physical and chemical properties inside cells, which in turn affect cellular function (Lippmann et al., 2019). Many enzymes lose their native folding that impairs their catalytic function, with disrupted activity at temperatures even slightly above their optimum (Daniel et al., 2008; Hobbs et al., 2013). These denatured or unfolded proteins can also disrupt protein homeostasis and form toxic insoluble aggregates (Richter et al., 2010; Guihur et al., 2022). High temperatures also increase membrane fluidity, altering its permeability to water, ions, and solutes, and destabilizing transmembrane proteins (Niu and Xiang, 2018; Cano-Ramirez et al., 2021). These changes in membrane stability can disrupt photosystem II (PSII) function and photosynthetic electron transport (Hugly et al., 1989). This can culminate in the accumulation of reactive oxygen species (ROS) that cause further damage to proteins, DNA, and lipids (Suzuki and Mittler, 2006). Therefore, heat can severely impact plant growth and fitness in the absence of appropriate acclimatory responses.

Molecular responses to heat have been extensively explored in model plants and crops, such as *Arabidopsis thaliana* and *Oryza sativa*. Such studies reveal that heat responses involve the regulation of heat-shock proteins (HSPs), which are predominantly chaperones and transcription factors (TFs) (Andrási et al., 2021; Guihur et al., 2022). The molecular chaperones, such as HSP70, HSP90, HSP100, and small HSPs, bind unfolded proteins to prevent their aggregation and, in some cases, induce their correct re-folding or degradation (Richter et al., 2010). The heat-shock transcription factors (HSFs) transcriptionally regulate an array of genes that can contribute to stress responses, including those encoding HSPs as well as others related to ROS signalling and scavenging, redox homeostasis, ion transport, metabolite biosynthesis (e.g. flavonoids), pathogen immunity, and growth and development (Richter et al., 2010; Andrási et al., 2021; Guihur et al., 2022). In addition, various phytohormones, such as abscisic acid (ABA) and ethylene, can influence heat stress response by regulating cellular processes including sucrose metabolism and ROS detoxification (Li et al., 2020a). This combinatorial response helps maintain cellular function under warmer temperatures (Yamori et al., 2014; de Pinto et al., 2015). Experimental heat-shocks, which often involve a temperature change of >10 ºC (Mittler et al., 2012), have been useful to identify the first responders to heat stress. However, plants also make metabolic and physiological adjustments to optimize their photosynthetic performance under more moderate (<10 ºC) and prolonged warming (Larkindale and Huang, 2004; Way and Yamori, 2014; Yamori et al., 2014). Such responses to longer-term warming may involve distinct pathways compared to acute responses (Wang et al., 2018; Wang et al., 2020), such as increasing the thermal optimum for photosynthesis, the electron transport rate, and/or stabilising RIBULOSE 1,5-BISPHOSPHATE CARBOXYLASE-OXYGENASE (RUBISCO) ACTIVASE (Way and Yamori, 2014; Yamori et al., 2014). Since global warming will likely cause gradual and sustained temperature increases, it is prudent to extend our understanding of thermal tolerance to moderate and sustained warming.

Here, we studied the transcriptomic response of *Wahlenbergia ceracea*, an herbaceous perennial that grows across Australia’s alpine regions, to prolonged moderate warming. We generated an annotated genome assembly and inferred functional attributes for thousands of genes based on homology to *A. thaliana*. Using this information, we contrasted changes in gene regulation in response to a daytime growth temperature difference of 6 °C, between heat-tolerant and heat-sensitive lines of *W. ceracea* (Arnold et al., 2024), which was shown to impact photosystem thermostability (Notarnicola et al., 2021; Arnold et al., 2022). This revealed differential warming-responsive regulation of genes involved in RNA regulation and photosynthesis. While only weak correlations could be made between co-expressed genes and thermal tolerance, stronger correlations could be established for chlorophyll content and maximum photosynthetic efficiency. The prediction of associated transcription factors suggested that this contrasting regulation may be related to phytohormone signaling pathways, especially ethylene and abscisic acid. These findings highlight candidate pathways that may contribute to heat tolerance in *W. ceracea* and underline the importance of expanding genomic investigation of native species.

## Materials and Methods

### Plant germplasm

*Wahlenbergia ceracea* Lothian (Campanulaceae) germplasm studied herein were derived from an established population (Notarnicola et al., 2021; Arnold et al., 2022; Arnold et al., 2024). Briefly, F_0_ plants were grown from seeds collected along an elevation gradient within the species’ subalpine-alpine distribution range (1590-2095 m a.s.l., Kosciuszko National Park, NSW, Australia). F_1_ plants were produced by crossing F_0_ individuals occupying similar elevations (< 50 m vertical elevation). The F_2_ generation was produced from F_1_ plants using a partial diallel half-sib breeding design to produce lines with controlled relatedness and systematic genetic variation. Subsequently, outcrossing was performed between unrelated F_2_ plants to produce the F_2_ seed that gave rise to the F_3_ plants used for temperature experiments herein. The pollen donor and recipient history for lines utilized in this study is provided in **Table S1**.

### Plant growth conditions and heat tolerance assessment

Seeds from F_2_ plants were germinated in seed raising mix in 14×8.5×5 cm pots in a glasshouse at 28/20 ºC (day/night) to maximise germination and establishment (Arnold et al., 2022; Notarnicola et al., 2023). Germinated seedlings were transplanted into individual pots (6.5×6.5×16 cm) containing seed raising mix supplemented with low-phosphorus slow-release fertilizer (Osmocote Plus, Scotts Australia) and grown in a temperature-controlled glasshouse. Plants were bottom watered as needed and supplemented with liquid fertiliser every two weeks (Yates Thrive, Yates Australia). Heat tolerance was assessed in glasshouse-grown F_3_ plants by measuring the temperature-dependent change in basal chlorophyll fluorescence to calculate *T*_*max*_ as described previously (Arnold et al., 2021).

### Temperature treatments

For temperature treatments, plants were grown in temperature-controlled growth capsules (Photon Systems Instruments, Brno, Czech Republic). Approximately three-month-old glasshouse-grown plants were moved into the growth capsules and allowed to acclimate for two weeks (28/20 ºC, 60% relative humidity, 14-hour photoperiod, 1200 µmol photons m^-2^ s^-1^). Subsequently, plants were moved to either cool [24/15 ºC day/night, representative of sub-alpine summer temperatures (Arnold et al., 2024)] or warm temperatures (30/20 ºC day/night). Plants grown at 30 ºC have reduced fitness (seed number and viability) but are neither sterile nor exhibit severe photoinhibition (Notarnicola et al., 2021). After two-weeks of growth, approximately 100 mg of tissue was collected from the youngest, fully-expanded leaves, which should have developed primarily under these contrasting temperatures. This experiment was performed in biological triplicate using full-sibling plants from the F_3_ lines defined as heat-tolerant and heat-sensitive. Replicates were randomly arranged across three blocks (one replicate per block), which was mirrored for the two treatments. Leaves were harvested in a randomized order between 11:00 and 16:30 hours, and immediately flash-frozen in liquid nitrogen and stored at -80 ºC. Time of harvesting and developmental stage were recorded for each sample (non-flowering n = 56; early bud n = 16).

### Flow cytometry

Flow cytometry was used to estimate genome size. Leaf material was excised from glasshouse-grown *W. ceracea* and kept on ice until processing. CyStain PI Absolute P (Sysmex Partex GmbH, Görlitz, Germany) was used for sample preparation following the manufacturer’s instructions, except for halved reaction volumes. Leaf material from soybean (*Glycine max* (L.) Merr. ‘Polanka’, 2C = 2.50 pg) (Doležel et al., 1994) was used as an internal standard. A tissue sample of approximately 0.5-1.0 cm^2^ was placed in 300 μL of Nuclei Extraction Buffer in a Petri dish and chopped manually with a clean razor blade. The lysate was filtered through a 40 μm cell strainer into a sample tube and mixed with 1,000 μL Staining Buffer supplemented with 50 μg propidium iodide and 50 μg RNAse. The sample was then loaded into a BD Accuri C6 Plus flow cytometer equipped with a 488 nm laser and a BD CSampler Plus (BD Biosciences, San Jose, CA, USA) and run at a flow rate of 14 mm min^-1^. Histogram data were collected using the FL2 detector while eliminating events with a value of less than 80,000 on FSC-H. Analysis was performed with the BD Accuri C6 Software (v1.0.23.1).

### DNA sequencing

High-molecular weight genomic DNA was extracted using an SDS-based protocol (Jones et al., 2021). Size selection was performed to obtain fragments ≥10 kb using the Short Read Eliminator (Circulomics) followed by isolating 12 kb fragments using a SageELF (Sage Science). A PacBio Sequel II long-read DNA sequencing library was prepared using the SMRTbell Express Template Prep Kit 2.0 (PacBio) and sequenced on one 8M SMRT cell, using the circular consensus sequencing mode to generate high-accuracy HiFi reads. Sequencing output and read quality was inspected with NanoPlot (v1.28.2) (De Coster et al., 2018).

### Ploidy analysis

Ploidy was assessed using k-mer count analysis on unassembled reads. K-mer frequency for k = 17, 21, 27, and 31 bp were determined using Jellyfish (Marçais and Kingsford, 2011). Coverage distributions at each k-mer length were analysed using Genomescope2.0 (Ranallo-Benavidez et al., 2020). Ploidy was also estimated with SmudgePlot (Ranallo-Benavidez et al., 2020) based on the 21-mer output from GenomeScope2.0.

### Genome assembly

PacBio HiFi long-reads were assembled using Canu v2.0 (Koren et al., 2017) with PacBio HiFi settings (Nurk et al., 2020) and a genome size estimate of 0.9 Gb. Contigs annotated as low confidence or representing alternate haplotypes were removed and assembly statistics were calculated using seqkit and quast (Gurevich et al., 2013; Shen et al., 2024). Reads were aligned against the assembly produced using minimap2 (Li, 2018), and read depth was calculated across 1 kb intervals using BEDTools makewindows (Quinlan and Hall, 2010). Repeats were identified using RepeatModeler (v2.0.7), and the resulting library was used to soft-mask the assembly with RepeatMasker (v4.2.2) (Flynn et al., 2020).

### mRNA sequencing

Total RNA was extracted from snap-frozen and ground leaf tissue using the Spectrum™ Plant Total RNA Kit (Merck, New Jersey, USA) with on-column DNase I digestion. Purified RNA was inspected for DNA contamination and integrity using semi-denaturing agarose-gel electrophoresis and the LabChip GX Touch (PerkinElmer). RNA was quantified with ND-1000 Spectrophotometer (NanoDrop Technologies) and Qubit 4 Fluorometer (RNA HS kit, Thermo Fisher Scientific). The cDNA libraries were constructed with 250 ng RNA using the Illumina stranded mRNA prep ligation protocol, as per manufacturer’s instructions except that reaction volumes were scaled by one-half and 12 PCR cycles were used for library amplification. Paired-end sequencing (2×150 bp) was conducted on the NovaSeq6000 (Deakin Genomics Centre, Deakin University, Victoria, Australia).

### Genome annotation

The soft-masked genome assembly was used for gene prediction with BRAKER2 (v2.1.5, --epmode --softmasking) (Stanke et al., 2006; Brůna et al., 2021). Gene predictions were guided by all eudicot orthogroups available in OrthoDB (v10) (Kriventseva et al., 2019) and mRNA sequencing generated herein (see below). For the latter, trimmed reads across all samples were merged and aligned to the genome using Subjunc (v1.6.2) (Liao et al., 2013). The amino acid and coding sequences were drawn from the braker.gtf output using “getAnnoFastaFromJoingenes.py” from Augustus.

Gene-space completeness was evaluated with Benchmarking Universal Single-Copy Orthologs (BUSCO, v6.0) based on embryophyta_odb10 and eudicot_odb10 lineages (Seppey et al., 2019). Functional information of predicted genes was inferred based on orthology between *W. ceracea* and *A. thaliana*. Orthologs were identified using OrthoFinder with default settings (Emms and Kelly, 2019) using protein sequences for the following species: *W. ceracea, A. thaliana, Eucalyptus grandis, Glycine max, Manihot esculenta, Oryza sativa, Solanum lycopersicum, Vitis vinifera, Zea mays, Chlamidomonas reinardtii, Marchantia polymorpha, Physcomitrium patens*, and *Selaginella moellendorfii*.

### Gene expression analyses

Raw reads were trimmed to remove adapter sequences and low-quality base calls (PHRED <20) using Trim Galore! with default settings. Trimmed reads were inspected using MultiQC (Ewels et al., 2016) prior to transcript quantification with Kallisto (Bray et al., 2016). The kallisto index was generated using coding sequences from the BRAKER2 predicted genes, which were extracted using GffRead (Pertea and Pertea, 2020). Only sequences with predicted exons, and start and stop codons were included. Transcript-level abundance estimates were summarised to gene-level counts with Tximport using the ‘length-scaled transcripts per million’ equation (Soneson et al., 2015). Only genes with >10 counts in at least 18 samples were retained for analysis. Counts were normalized using the Trimmed Mean of M-values (Robinson and Oshlack, 2010). Observation-level and sample-specific quality weights were estimated based on mean-variance modelling using voomWithQualityWeights (limma) to account for heteroscedasticity (Liu et al., 2015). To account for samples derived from the same plant line, we estimated within-line correlation using duplicateCorrelation (limma) which was then incorporated into subsequent linear models (Ritchie et al., 2015). Linear modelling was performed with lmFit (limma) using gene-level log-CPM values:

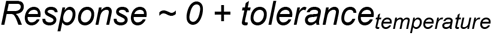

We utilized a pseudo-factor (*tolerance*_*temperature*_) that combines tolerance group and growth temperature (Tolerant_cool_, Sensitive_cool_, Tolerant_warm_, and Sensitive_warm_; n per group = 18). The no-intercept model allowed for direct comparisons in expression levels between sample groups. Enriched functional terms were identified using the deduplicated *A. thaliana* orthologs of predicted *W. ceracea* genes with Metascape (Zhou et al., 2019). Enrichment analyses were based on GO Biological Process, KEGG Pathway, and WikiPathway catalogues with default settings (minimum overlap = 3; P-value cut-off = 0.01; minimum enrichment factor = 1.5). TFs were associated with *A. thaliana* orthologs of up- and down-regulated genes, separately, for tolerant and sensitive *W. ceracea* lines using TF2Network (Kulkarni et al., 2018). Co-expressed gene networks were constructed via weighted gene co-expression network analysis (WGCNA) (Langfelder and Horvath, 2008) using log2-transformed TMM-normalized counts averaged across tolerance_temperature_ groups. A soft-thresholding power was determined to satisfy scale-free topology (14, R^2^ >0.9), which was used to build an unsigned adjacency matrix. This was converted into an unsigned Topological Overlap Matrix as a measure of similarity. Subsequently, gene modules were created with dynamic tree cut and module eigengenes were calculated. Module eigengenes were used for correlating gene expression patterns to physiological trait variation. For this analysis, we utilized previously reported trait measurements that were taken on sibling plants derived from the same parental crosses as studied here: critical thermal limits of PSII (T_crit-hot_, T_crit-cold_), chlorophyll content, maximum quantum efficiency of PSII (F_v_/F_m_), and reproductive stem count (Arnold et al., 2024). Since these measurements were made on different individuals, module eigengenes were correlated to mean trait values from measurements taken across 5-8 plants from each tolerance_temperature_ group.

## Results

### Genome assembly and annotation

We performed *de novo* genome assembly for *W. ceracea* using PacBio HiFi long-read sequencing. This was performed on genomic DNA from a single F_2_ individual (F_2_.209, **Table S1**) from an established *W. ceracea* population (Notarnicola et al., 2021; Arnold et al., 2022; Arnold et al., 2024). This individual represents a mixed pedigree derived from relatively low-elevation (F_0_.1613×F_0_.1606, ∼1,750 m) and high-elevation (F_0_.1525×F_0_.1524, ∼1,900 m) F_0_ founders, which should reflect the genetic variation across the sampled elevation gradient and be present across the F_3_ individuals studied herein. After removing low-quality (PHRED <20), low-coverage, and non-plant reads, we retained ∼1.8 million reads (mean length=12.2 kb, mean Phred score = 33.5). Ploidy analysis using GenomeScope and Smudgeplot on unassembled reads (Ranallo-Benavidez et al., 2020) supported a diploid genome (**Figure S1**), and flow cytometry estimated a nuclear DNA content of ∼1.8 Gb. Using the filtered HiFi reads with haploid genome size estimation of 0.9 Gb, Canu produced a 1.37 Gb assembly (∼76% of the expected diploid genome size) with an estimated 25× sequencing coverage across 4,820 contigs (**Figure 1A**, N_50_ = 834 kb, L_50_ = 447, GC content = 39.71%). We observed that removing repeat-rich contigs reduced the assembly size to ∼810 Mb and increased contiguity (2,047 contigs, N_50_ = 1.12 Mb, L_50_ = 210). Employing RepeatModeler and RepeatMasker (Flynn et al., 2020) revealed that ∼72% of our assembly was composed of repeats, with Long Terminal Repeat (LTR) retroelements being the most abundant class present (∼40%; **Table S2**). BUSCO analyses using the Embryophyta and Eudicot lineages showed high gene-space completeness for the genome (**Figure 1B**). However, there was a large proportion of duplicated BUSCOs in the raw assembly, which decreased after stepwise removal of junk contigs, putative haplotigs, and repeat-annotated contigs. This is consistent with a raw assembly retaining uncollapsed haplotypes and repeat-rich regions that inflate BUSCO duplication.

**Figure 1.**
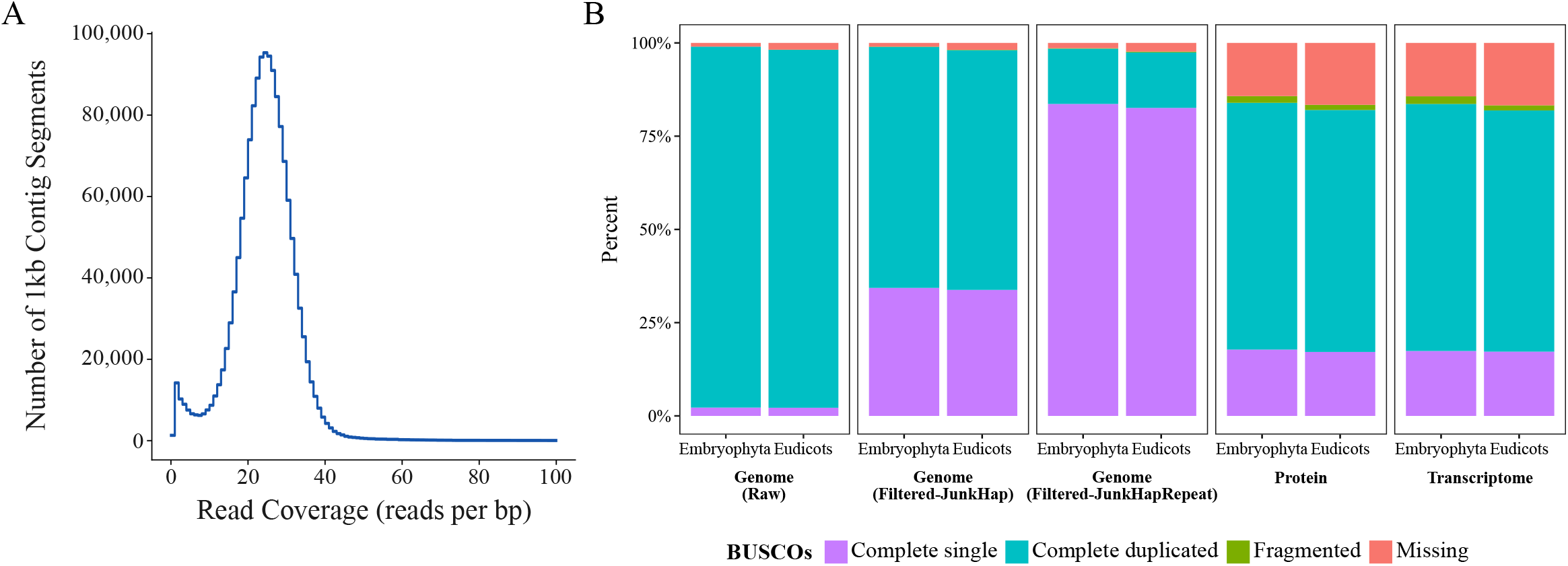
*W. ceracea* genome assembly. (A)Histogram of per-base read coverage binned into 1 kb windows along contigs of the genome assembly. (B)BUSCO completeness, based on embryophyte and eudicot databases, for the genome assembly, protein sequence (Protein), and coding sequence annotations (Transcriptome). Three versions of the genome assembly were (1) an unfiltered assembly (Raw), (2) an assembly with junk contigs and haplotigs removed (Filtered- and (3) an assembly with junk contigs, haplotigs, and repeat-annotated contigs removed (Filtered-JunkHapRepeat).

After soft-masking with RepeatMasker, we annotated the genome using BRAKER2 (Brůna et al., 2021) augmented with the mRNA sequencing performed herein. This yielded 223,827 exons across 87,730 predicted protein-coding genes. BUSCO assessment of the BRAKER2 protein and transcript annotations showed reduced completeness compared to the genome assembly, likely due to limited transcript support for genes not expressed in leaf tissue (**Figure 1B**). Nevertheless, using these annotated coding sequences, we performed transcript quantification per sample with Kallisto (Bray et al., 2016). On average, ∼69.7% of reads could be assigned to an annotated transcript (**Supplementary Data 1**). In total, we captured 38,257 expressed transcripts that summarised to 37,307 genes, which is comparable to numbers reported in *A. thaliana* (48,359 transcripts; 27,655 genes) (Cheng et al., 2017). For functional annotations, we used OrthoFinder (Emms and Kelly, 2019) to infer orthologous genes across a range of plant species. *W. ceracea* shared the highest number of orthogroups with *Manihot esculenta* (9,357), *Glycine max* (9,275), and *Vitis vinifera* (9,239) among the species analysed (**Table S3**). Importantly, there were 11,205 orthologs between *W. ceracea* and *A. thaliana*, from which we could derive functional information (**Supplementary Data 2**). Overall, we identified *A. thaliana* orthologs for ∼81.4% warming-responsive DEGs in *W. ceracea* (see below). Therefore, this genome assembly and the associated gene annotations provide a robust basis for investigating warming-responsive gene regulation in *W. ceracea*.

### Contrasting regulation of RNA metabolism and photosynthesis-associated genes between heat-tolerant and heat-sensitive lines of *W. ceracea*

We inferred thermal tolerance based on the thermal limit of photosynthesis as assessed by chlorophyll fluorometry (Geange et al., 2021). More specifically, we measured the temperature-dependent change in basal chlorophyll fluorescence to derive *T*_*max*_ (the temperature at maximum fluorescence), which is posited to represent the point of irreversible PSII damage (Berry and Bjorkman, 1980; Knight and Ackerly, 2002; Arnold et al., 2021; Posch et al., 2026) across 24 F_3_ lines (**Figure 2A**).

**Figure 2.**
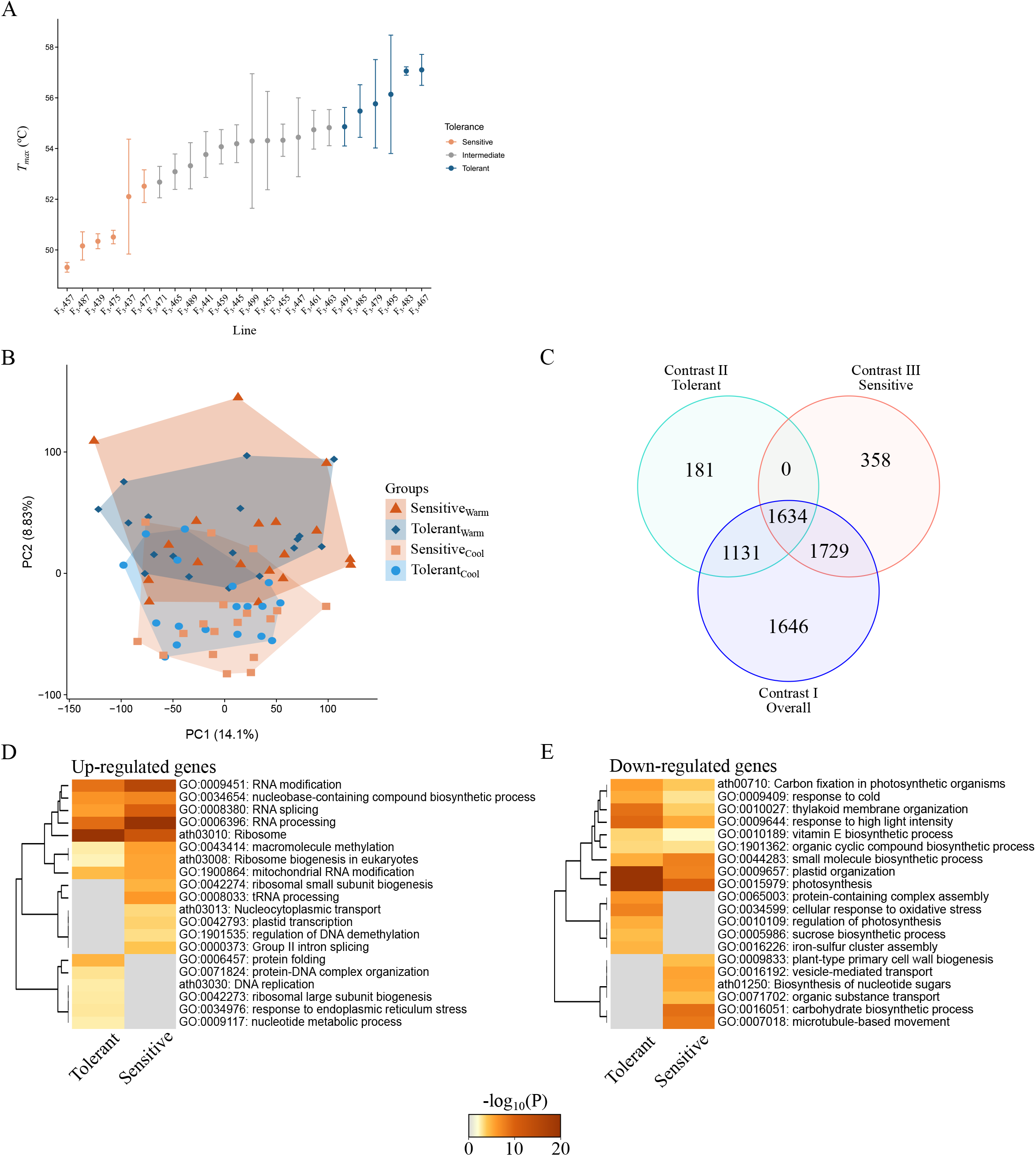
Temperature-responsive gene regulation in heat-tolerant and heat-sensitive lines of *W. ceracea*. (A)Measures of *T*_max_ across individuals from 24 F_3_ lines of W. ceracea. Points denote means (n=2-3) and error bars denote standard error of the mean. (B) PCA on gene-level expression of detected genes across all samples. (C) Venn diagram of heat-responsive DEGs detected in each contrast defined in Table 1. (D-E) Heatmaps with one-dimensional hierarchical clustering of significantly enriched GO Processes and KEGG Pathway terms in (D) up- and (E) down-regulated genes in heat-tolerant and heat-sensitive lines. Significant terms were clustered by Metascape and the most significant terms are shown. Cell colour denotes statistical significance (-log_10_ P-value; grey presents non-significant terms).

Subsequently, these lines were ranked by mean *T*_*max*_ to identify those with the six highest and lowest values, which were defined as relatively heat-tolerant (mean *T*_*max*_ = 56.1 °C) and heat-sensitive (mean *T*_*max*_ = 50.8 °C; Δ*T*_*max*_ = 5.3 °C), respectively. These categorized lines formed the basis for exploring gene expression patterns associated with thermal tolerance.

We then investigated warming-responsive gene regulation, associated with differences in thermal tolerance, by performing mRNA sequencing on six individuals from each line (i.e. 36 relatively heat-tolerant and heat-sensitive individuals) exposed to two weeks at either cool (24/15 ºC) or warm (30/20 ºC) growth conditions. In total, we reliably detected (>10 counts in ≥18 samples) 22,585 of the BRAKER2-predicted genes.

Principal Component Analysis (PCA) revealed separation along PC2 (8.83%), which reflected growth conditions (**Figure 2B**). PC1 (14.10%) did not associate with any measured variable, including sampling time, and therefore possibly reflects genetic differences. Next, we confirmed that all individuals were responsive to warmer conditions through the detection of 6,140 warming-induced DEGs when incorporating all samples (Contrast *I*, **Table 1, Figure 2C**). Among up-regulated genes, we observed an enrichment of those associated with RNA processes, including splicing, modification, and translation (i.e. ribosome), whereas down-regulated genes were related to photosynthesis, light harvesting, and plastid organization (**Figure S2A, Supplementary Data 3**). Interestingly, there was a lack of heat stress-related terms, which is consistent with the hypothesis that moderate warming elicits a different response to heat-shock (Guihur et al., 2022).

**Table 1:**
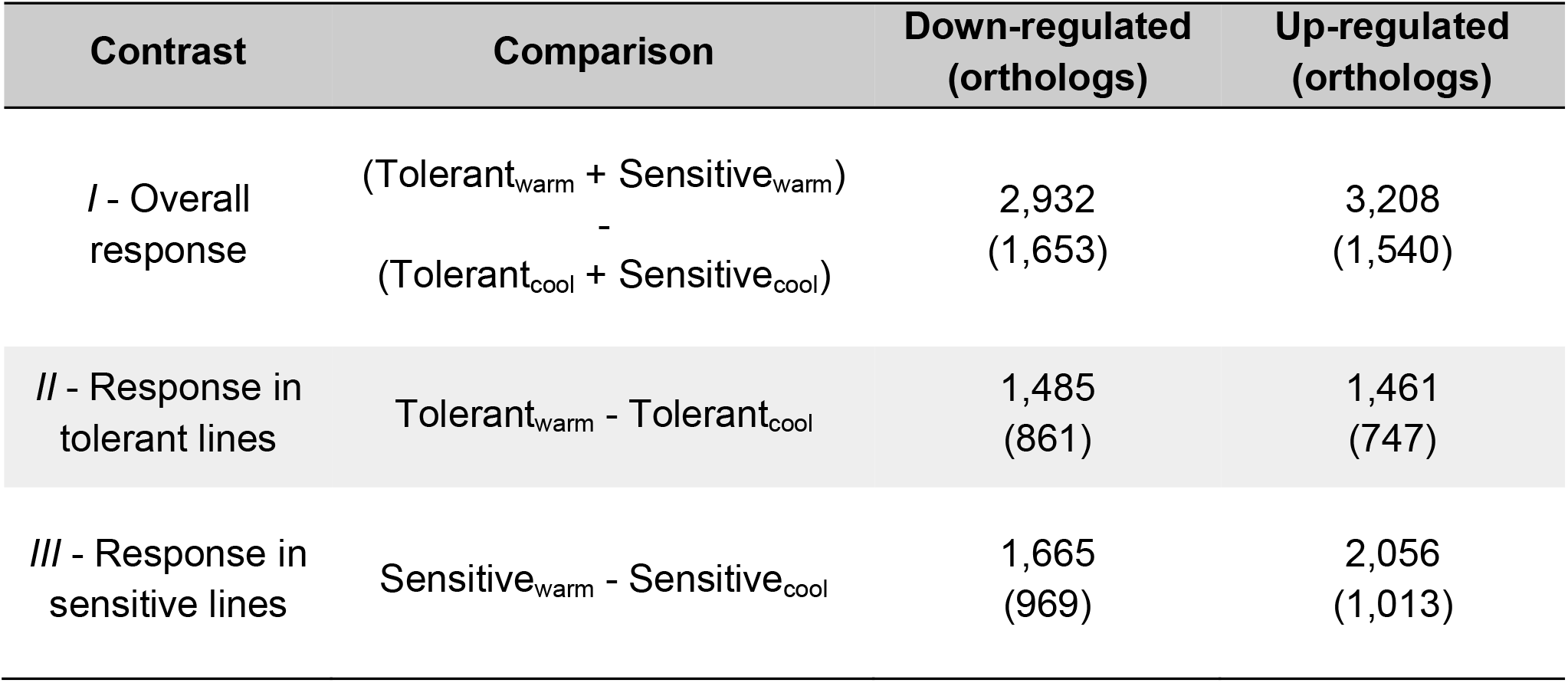
Number of temperature-responsive genes identified from each contrast performed in *W. ceracea*.

We then contrasted warming-induced DEGs between heat-tolerant (Contrast *II*) and heat-sensitive lines (Contrast *III*). There were 2,946 and 3,721 DEGs in tolerant and sensitive lines, respectively (**Figure 2C, Table 1, Supplementary Data 4-5**). Interestingly, these comparisons missed 1,646 DEGs that were significant in Contrast *I*, likely the result of reduced statistical power with smaller sample size. Nonetheless, both tolerant and sensitive lines showed pronounced warming-induced changes in expression (**Figure S2B**). We contrasted enriched functional terms found amongst warming-responsive genes between tolerant and sensitive lines (**Figure 2D-E, Supplementary Data 6**). Although we observed similar functional terms identified in Contrast *I*, this analysis revealed that heat-sensitive lines had a more pronounced enrichment for up-regulating RNA processing-related genes than heat-tolerant lines. On the other hand, heat-tolerant lines had a stronger enrichment for down-regulated genes related to photosynthesis. We further characterized the changes in expression of photosynthesis-associated genes that were down-regulated under warm conditions in heat-tolerant lines (**Figure S3**). Although these genes were down-regulated in both lines, tolerant lines demonstrated a stronger down-regulation, especially for a subset of genes that included *LIGHT HARVESTING COMPLEXES (LHCs), OXYGEN EVOLVING COMPLEX 33* (*OEC33*), and *PHOTOSYNTHETIC ELECTRON TRANSPORT C (PETC)*. Together, these results suggest that heat-tolerant and heat-sensitive lines exhibit contrasting modulation of RNA regulation and photosynthesis-associated genes during warming.

### Co-expression analysis reveals gene-trait correlations for chlorophyll content and photosynthetic efficiency, but not heat tolerance, during moderate warming

We employed WGCNA (Zhang and Horvath, 2005; Langfelder and Horvath, 2008) to correlate expression patterns of specific genes to variation in T_max_ alongside additional traits, associated with thermal tolerance and photosynthesis, that were measured in sibling plants (Arnold et al., 2024). In total, seven co-expressed gene modules were identified and their expression patterns were summarized using eigengenes, which were then correlated to mean trait values obtained from heat-tolerant and heat-sensitive plants (**Figure 3A, Supplementary Data 7**). Despite our sampling strategy, we did not observe any modules that significantly correlated with T_max_.

**Figure 3.**
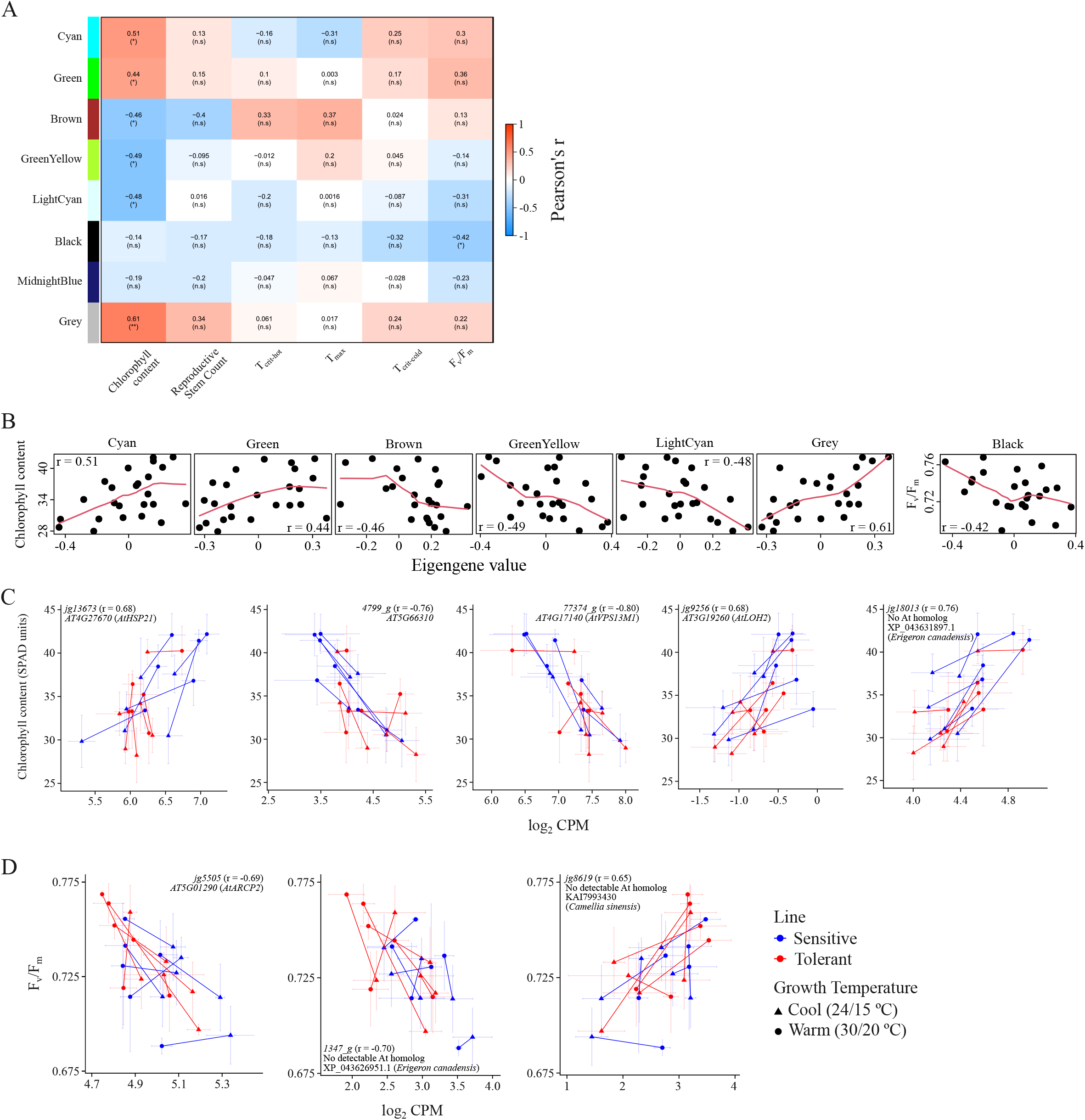
Co-expressed gene networks reveal module- and gene-trait relationships in heat-tolerant and sensitive lines of *W. ceracea*. (A)Heatmap representing the correlation (Pearson’s r) between module eigengene values and mean trait values measured in sibling plants from F_3_ *W.ceracea* lines. Student asymptotic p-values were calculated to determine statistically significant correlations: n.s., not significant; ^*^, p < 0.05; ^**^ p < 0.01. (B) Scatterplots between trait values and module eigengenes. Pearson’s r values are shown as a measure of strength and direction for each relationship. (C-D) Scatterplots showing the relationship between mean trait values and mRNA abundance (log CPM) in heattolerant and sensitive lines of *W. ceracea* grown, in warm and cool conditions, for select gene-trait relationships for (C) chlorophyll content and (D) F_v_/F_m_. Gene significance (Pearson’s r), BRAKER2-generated gene ID, and detected Arabidopsis homolog is shown for each gene-trait plot (or NCBI sequence ID and species for the most similar homolog as determined by protein BLAST).

However, we did observe significant module-trait correlations with chlorophyll content and F_v_/F_m_ (**Figure 3B**). We further interrogated these modules for gene-trait correlations based on gene significance scores (correlation of gene-level abundance and trait values, **Supplementary Data 8**). In each modules, we could observe strong gene-trait correlations (|Pearson’s r| >0.6) in both directions, likely the result of using unsigned networks in our co-expression analysis (**Figure 3C-D**). Nevertheless, these included many genes with detectable *A. thaliana* homologs, which implicated proteins of diverse functions ranging from stress response (e.g. *AT4G27670*), lipid metabolism (e.g. *AT3G19260*), and RNA processing (e.g. *AT5G01290*). However, there were also many well-correlated gene-trait relationships for genes without a detectable *A. thaliana* homolog. For instance, a BLAST of the predicted protein sequence for *jq18103, 1347_g*, and *jg8619*, returned homologs from species within asterids (containing *W. ceracea*) but not Brassicaceae (containing *A. thaliana*). This suggests that we have identified lineage-specific temperature-responsive genes that could contribute to heat acclimation.

### Heat-sensitive *W. ceracea* repress gene networks related to development and reproduction under warm conditions

Since we observed contrasting warming-responsive regulation between heat-tolerant and heat-sensitive lines, we identified candidate TFs that might explain these differences. Using TF2Network to construct gene regulatory networks (Kulkarni et al., 2018), we identified a total of 23 and 396 TFs based on up- and down-regulated genes, respectively (**Figure 4A-B, Supplementary Data 9**). Interestingly, most TFs were associated with down-regulated genes, with 381 and 98 detected in sensitive and tolerant lines, respectively, of which 83 were in common. This bias suggests that TF inactivation is more pronounced during warming, and that this is more pronounced in heat-sensitive lines. We found that the sensitive-specific TFs targeting down-regulated genes were related to biosynthetic and developmental processes, as well as hormone signaling, especially ethylene-activated signalling, indicating that sensitive lines are responding differently to growth and developmental cues during warming (**Figure 4C**). Among those TFs identified targeting down-regulated genes in both lines, we observed enrichments related to ABA signaling and light stress, highlighting that both lines are responsive to these environmental cues. We then investigated whether we observed a bias towards warming-induced expression regulation of the genes encoding the identified TFs. Only 45 TF-encoding genes were differentially expressed, of which 26 were down-regulated, including multiple TFs responsive to signalling molecules such as ABA and ethylene (**Figure 5**). Although these general trends were observed, numerous TF-encoding genes also showed distinct steady-state levels under cool and warm conditions between lines (e.g. *41073g, 62467g, jg2041*). Interestingly, only a handful of TF-encoding genes demonstrated contrasting regulation between lines, including *jg9195, 77210g*, and *g8224*. Therefore, we have identified a range of TFs that may contribute to warming-induced gene regulation in *W. ceracea*, including specific TF-encoding genes that demonstrate contrasting regulation between heat-tolerant and heat-sensitive lines.

**Figure 4.**
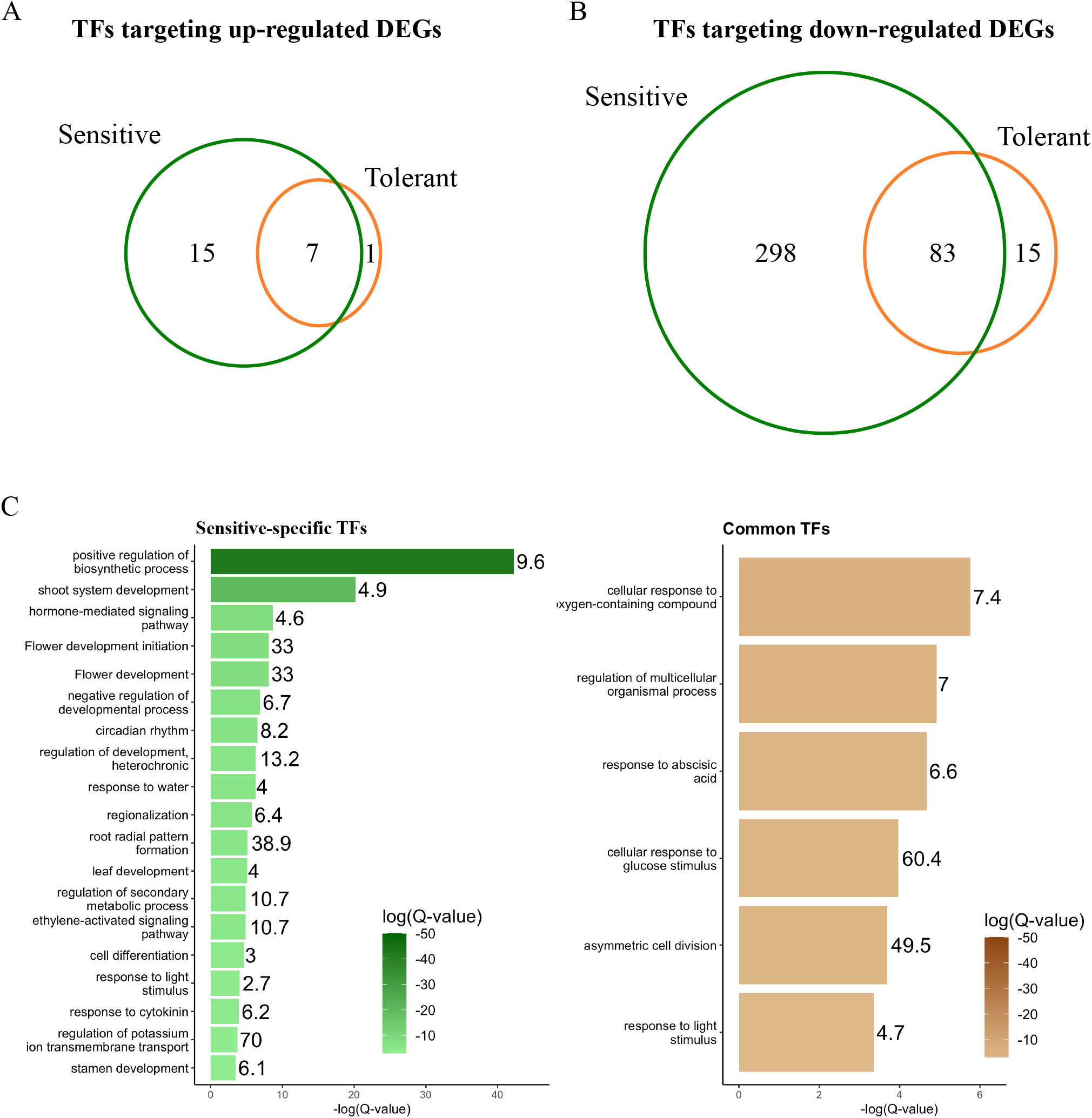
Transcription factors predicted to target heat-responsive genes. (A-B) Venn diagrams of the number of TFs identified by constructing gene regulatory networks with *TF2Network* using (A) up-regulated and (B) down-regulated genes detected in heat-tolerant and sensitive lines of *W. ceracea*. (C) Bar charts representing enriched functional terms for TFs, based on detected Arabidopsis orthologs, identified as targeting down-regulated genes in sensitive (sensitive-specific TFs) or both (common TFs) lines. Numbers denote the fold enrichment of genes in each category.

**Figure 5.**
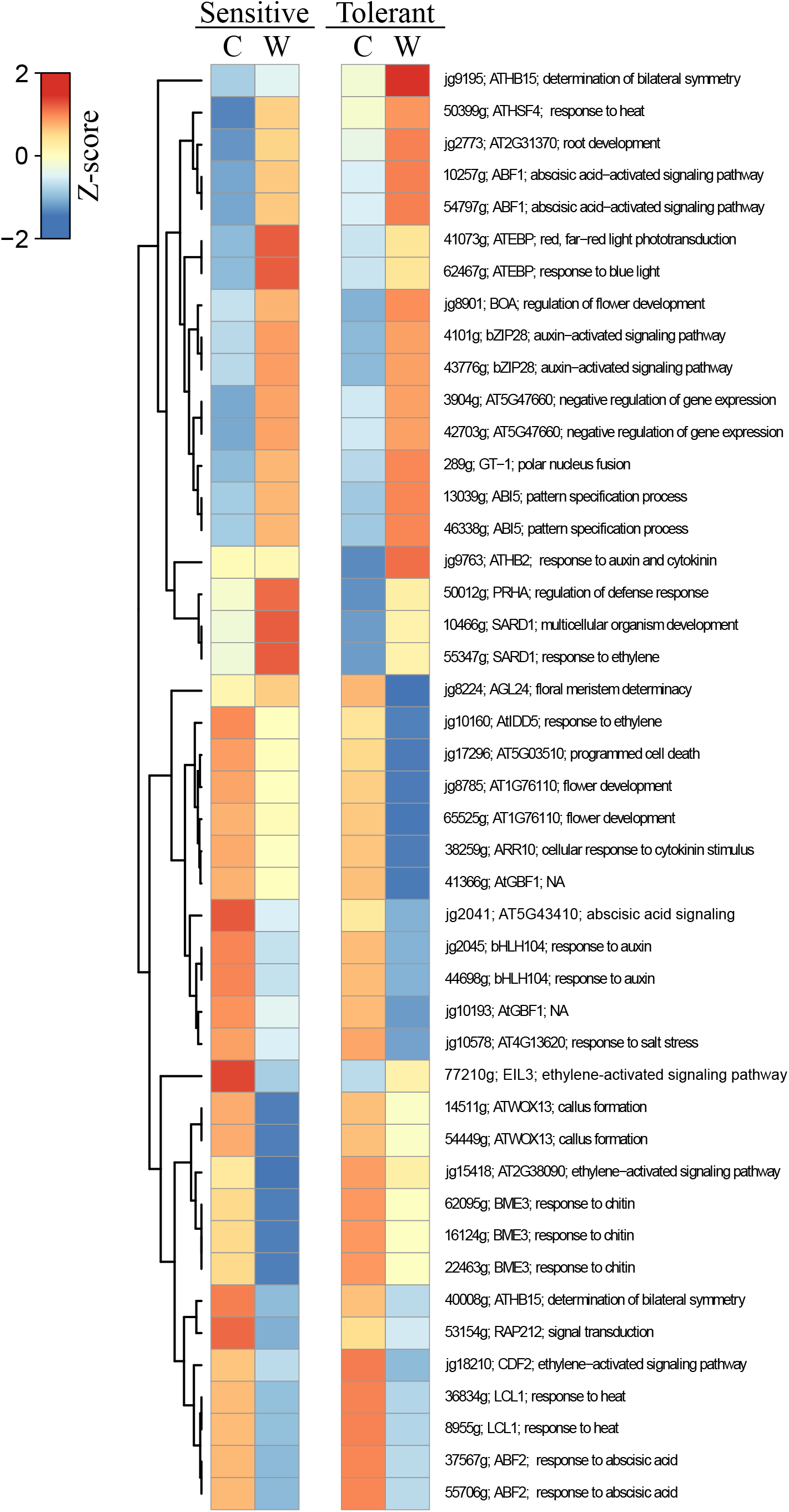
Differentially expressed transcription factors associated with temperature-responsive gene networks in *W. ceracea*. Heat map with one-dimensional hierarchical clustering representing mean expression (Z-scores) of genes encoding transcription factors, identified by TF2Network, which are differentially expressed between cool (C) and warm (W) conditions. Expression across *W. ceracea* paralogs was averaged. Labels denote *W. ceracea* gene ID, Arabidopsis gene symbol, and the most significant GO process.

## Discussion

Elucidating how wild plants respond to warmer temperatures could improve our ability to predict the impacts of climate change. Here, we produced an annotated genome assembly for the Australian alpine herb *W. ceracea*. With this, we investigated gene regulation during prolonged moderate warming in heat-tolerant and heat-sensitive lines. We identified thousands of warming-responsive genes, including hundreds that were differentially regulated between tolerant and sensitive lines. Specifically, we found that tolerant lines more prominently down-regulated genes associated with photosynthesis while up-regulating those related to translation and protein folding.

Conversely, sensitive lines up-regulated genes related to RNA modification and processing, and down-regulated those related to carbohydrate biosynthesis. Our findings also highlight the importance of studying longer-term warming, which appears to elicit a distinct response compared to heat-shock, in a native species whose genome may not be generalizable from model systems.

Our *W. ceracea* genome assembly had high gene-space completeness (>90%), as determined by BUSCO assessment (**Figure 1B**), but limited contiguity. This fragmentation is consistent for a large, heterozygous plant genome and likely reflects the sequencing of an outcrossed F_2_ individual. However, elevated heterozygosity is also expected in *Wahlenbergia* species, including *W. ceracea*, which exhibit a mixed-mating system characterised by predominant outcrossing, driven by protandry and herkogamy, and facultative autogamy (Anderson et al., 2000; Nicotra et al., 2015). We also observed substantial repeat content (∼72% of assembly), particularly of LTR retroelements (**Table S2**), which likely further impaired the ability to collapse haplotypes. While future investigations would benefit from sequencing inbred lines, to resolve haplotypes, or sampling wild accessions, to explore natural genetic variation, our assembly was sufficient for identifying protein-coding regions expressed during warming. Furthermore, the transposon-rich nature of the genome highlights *W. ceracea* as a promising system for studying the ecological and evolutionary impacts of transposon biology (Rey et al., 2016).

The temperature at maximum basal chlorophyll fluorescence (T_max_) reflects the thermal threshold for photosynthesis (Schreiber and Berry, 1977; Ilík et al., 2003; Ducruet et al., 2007). Therefore, a higher T_max_ should reflect an ability to sustain photosynthesis at higher temperatures (Berry and Bjorkman, 1980). However, the genes that determine T_max_ are yet to be identified and our mRNA-based analysis could only reveal weak associations. Although we had a mean Δ*T*_*max*_ = 5.3 °C between heat-tolerant and heat-sensitive lines (**Figure 2A**), it is possible that this between-group difference was insufficient, relative to intra-group variation, to produce statistically reliable gene-trait correlations. Indeed, greater variation in *T*_*max*_ can be observed between species (Berry and Bjorkman, 1980; Knight and Ackerly, 2002), positing that greater statistical power could be obtained from interspecies comparisons.

Nevertheless, utilizing the phenotypic plasticity we previously observed in these lines (Arnold et al., 2024), we could statistically establish positive and negative gene-trait relationships with warming-responsive changes in chlorophyll content and photosynthetic efficiency (**Figure 3C-D**), which might contribute to maintaining photosynthesis at warmer temperatures. While some of these genes appeared to encode homologs of well-characterized proteins in *A. thaliana* (e.g. *AtHSP21*), others only returned homologs from species outside of Brassicaceae. These could represent lineage-specific genes that will be enticing prospects for future research.

Heat-tolerant lines (higher T_max_) appeared to more strongly repress photosynthesis-related genes during warming (**Figure 2E**), including PSII components (e.g. *OEC33, ATPC1*) and antennae complexes (*LHCAs* and *LHCBs*, **Figure S3**), suggesting that transcriptional regulation of photosynthesis may contribute to thermal tolerance in *W. ceracea*. Similar regulation was also observed in *Nothofagus pumilio* where photosynthesis-associated genes were down-regulated in response to heat, including those involved in light harvesting and chlorophyll biosynthesis (Estravis-Barcala et al., 2021). How the repression of photosynthesis contributes to heat tolerance remains to be verified. One possibility is that reducing photosynthetic electron transport from PSII avoids PSI photoinhibition under prolonged warming in *W. ceracea*, which could otherwise cause redox imbalance and impaired photochemistry (Zivcak et al., 2014; Jiang et al., 2021). We also observed differential regulation of genes associated with RNA modification, processing, transport, and translation (**Figure 2D**).

Similar genes, related to RNA splicing and translation, were also induced by heat in *Nothofagus pumilio, A. thaliana*, and *Populus tomentosa* (Estravis-Barcala et al., 2021). These results are consistent with work highlighting a relationship between RNA regulation and stress responses. For example, heat and salinity reportedly caused changes in secondary structure and covalent modifications, which alter transcript stability (Anderson et al., 2018; Kramer et al., 2020). Heat-induced alternative splicing could produce protein isoforms with altered biochemical properties such as thermostability (Reddy et al., 2013), which has been observed for *RUBISCO ACTIVASE* in *Triticum aestivum* (Scafaro et al., 2019) and *HSFA2* in *A. thaliana* (Liu et al., 2013). Our observations suggest there may be transcriptional regulation of these processes during warming, which require follow-up via direct measurement of transcript half-lives, isoform switching, and RNA modifications to establish their potential ecological relevance.

We hypothesized that heat-tolerant and heat-sensitive lines would exhibit contrasting regulation of HSPs, especially molecular chaperones. Indeed, we observed that genes related to protein folding were up-regulated in heat-tolerant lines only (**Figure 2D**). This included many genes encoding chaperones in addition to HSPs, such as *BINDING PROTEIN 1* (or *HSP70-11*), *CALNEXIN 1*, and *CALRETICULIN 1A* (**Supplementary Data 4**), which are involved in stabilizing nascent proteins (Sun et al., 2021). This suggests an attempt to maintain protein homeostasis (proteostasis), which may include photosynthesis-associated proteins that are prone to destabilization during stress (Li et al., 2022). Further investigation of photosynthetic activity and protein stability in warm-grown *W. ceracea* would help determine the functional consequences of the gene regulation observed here. Nonetheless, these results suggest that tolerant lines repress photosynthesis and prioritize protein fidelity under warmer temperatures.

We associated hundreds of TFs with down-regulated genes in heat-sensitive lines (**Figure 4B**). These were enriched for functions related to biosynthetic pathways, development, and hormone signaling (**Figure 4C**). Those associated with biosynthetic and developmental pathways likely reflect regulation occurring as a consequence of increased heat sensitivity. However, the identification of homologs of *ETHYLENE-INSENSITIVE-LIKE 3* (*EIL3*) and multiple *ETHYLENE RESPONSE FACTORs* (*ERFs*) might reflect a contribution of ethylene signalling towards heat tolerance (**Supplementary Data 9**). Indeed, ethylene signaling, through *EIL3* and multiple *ERFs*, can stimulate heat responses by inducing *HSFs* (Huang et al., 2021; Poór et al., 2022). Furthermore, treatment of rice seedlings with the ethylene precursor, 1-aminocyclopropane-1-carboxylic acid, lead to increased chlorophyll *a*, up-regulation of *HSFs*, and increased detoxification activity that correlated with greater heat tolerance (Wu and Yang, 2019). Consistent with this, we observed that a homolog of *EIL3* was up-regulated during warming in heat-tolerant lines, but down-regulated in heat-sensitive lines (**Figure 5**). Therefore, ethylene signalling through *WcEIL3*, and other ERF homologs, represent promising candidates for contributing to heat tolerance in *W. ceracea*.

Contrary to expectations, we could only associate two HSFs, HSFC1 and HSF4, to warming-induced gene regulation. However, we could link 13 BASIC LEUCINE ZIPPER (bZIP) TFs to down-regulated genes (**Supplementary Data 9**), suggesting that they are more relevant during longer term moderate warming. This is supported by observations in *A. thaliana* where *HSFs* were regulated after short-term heat-shock but *bZIPs* were up-regulated after prolonged warming (Wang et al., 2020). The bZIP homologs identified here show evidence of contribution to heat tolerance in other species. For example, bZIP28 and bZIP60 promote the heat-shock response in maize (Li et al., 2020b) and *A. thaliana* (Kataoka et al., 2017), respectively. Only 45 of the 400 TF-encoding genes potentially regulating gene networks in our study were transcriptionally regulated by warming (**Figure 5**). Therefore future investigations should also consider whether post-translational modifications, like phosphorylation, rather than expression might influence TF activity (Richter et al., 2010; Andrási et al., 2021).

In conclusion, we generated an annotated genome assembly for *W. ceracea*, which we use to identify thousands of warming-regulated genes in heat-tolerant and heat-sensitive lines. From this, we propose that the warming-responsive regulation of genes associated with RNA metabolism, photosynthetic machinery, and protein chaperones, possibly coordinated through activity of hormone-responsive gene networks like ethylene, are important for thermal tolerance in this Australian native plant.

## Acknowledgements

This project was supported by the Australian Plant Phenomics Facility and the National Computational Infrastructure, both supported under the National Collaborative Research Infrastructure Strategy of the Australian Government. We thank the plant services team at The Australian National University for assistance with plant maintenance. We acknowledge the provision of expertise and resources from the Ecogenomics and Bioinformatics Lab, hosted within the Centre for Biodiversity Analysis at The Australian National University. We thank Tenzin Norzin, for technical assistance and plant maintenance, and Verónica Briceño and Marcin Adamski for insightful discussions.

## Author Contributions

RFN, PAA, ABN, and DRG designed experiments. RFN and PAA grew plants. PAA assessed thermal tolerance. RFN, ABN, and PAA harvested leaf samples. ANS performed flow cytometry. JH, AWJ, and BS conducted DNA sequencing and genome assembly. RFN and DRG performed RNA extractions. SL performed mRNA sequencing. RFN, DRG, and ZF conducted bioinformatic analyses. RFN, JH, and DRG wrote the manuscript. ABN and PAA edited the manuscript. All authors commented on the manuscript.

## Data availability

Genome and mRNA sequencing data have been deposited at NCBI (GenBank accession JBVYUB000000000, GEO accession GSE274951, BioProject accession PRJNA1148632). Gene and repeat annotations are available at Figshare (10.6084/m9.figshare.27890484).

## References

Anderson GJ, Bernardello G, Lopez PS, Crawford DJ, Stuessy TF (2000) Reproductive biology ofWahlenbergia (Campanulaceae) endemic to Robinson Crusoe Island (Chile). Plant Syst Evol 223: 109–123

Anderson SJ, Kramer MC, Gosai SJ, Yu X, Vandivier LE, Nelson ADL, Anderson ZD, Beilstein MA, Fray RG, Lyons E, et al (2018) N-Methyladenosine Inhibits Local Ribonucleolytic Cleavage to Stabilize mRNAs in Arabidopsis. Cell Rep 25: 1146–1157.e3

Andrási N, Pettkó-Szandtner A, Szabados L (2021) Diversity of plant heat shock factors: regulation, interactions, and functions. J Exp Bot 72: 1558–1575

Arnold PA, Briceño VF, Gowland KM, Catling AA, Bravo LA, Nicotra AB (2021) A highthroughput method for measuring critical thermal limits of leaves by chlorophyll imaging fluorescence. Funct Plant Biol 48: 634–646

Arnold PA, Wang S, Catling AA, Kruuk LEB, Nicotra AB (2022) Patterns of phenotypic plasticity along a thermal gradient differ by trait type in an alpine plant. Functional Ecology 36: 2412–2428

Arnold PA, Wang S, Notarnicola RF, Nicotra AB, Kruuk LEB (2024) Testing the evolutionary potential of an alpine plant: phenotypic plasticity in response to growth temperature outweighs parental environmental effects and other genetic causes of variation. J Exp Bot 75: 5971–5988

Berry J, Bjorkman O (1980) Photosynthetic response and adaptation to temperature in higher plants. Annu Rev Plant Physiol 31: 491–543

Bray NL, Pimentel H, Melsted P, Pachter L (2016) Near-optimal probabilistic RNA-seq quantification. Nat Biotechnol 34: 525–527

Brůna T, Hoff KJ, Lomsadze A, Stanke M, Borodovsky M (2021) BRAKER2: automatic eukaryotic genome annotation with GeneMark-EP+ and AUGUSTUS supported by a protein database. NAR Genom Bioinform 3: lqaa108

Cano-Ramirez DL, Carmona-Salazar L, Morales-Cedillo F, Ramírez-Salcedo J, Cahoon EB, Gavilanes-Ruíz M (2021) Plasma Membrane Fluidity: An Environment Thermal Detector in Plants. Cells 10: 2778

Cheng C-Y, Krishnakumar V, Chan AP, Thibaud-Nissen F, Schobel S, Town CD (2017) Araport11: a complete reannotation of the Arabidopsis thaliana reference genome. Plant J 89: 789–804

Daniel RM, Danson MJ, Eisenthal R, Lee CK, Peterson ME (2008) The effect of temperature on enzyme activity: new insights and their implications. Extremophiles 12: 51–59

De Coster W, D’Hert S, Schultz DT, Cruts M, Van Broeckhoven C (2018) NanoPack: visualizing and processing long-read sequencing data. Bioinformatics 34: 2666–2669

Doležel J, Doleželová M, Novák FJ (1994) Flow cytometric estimation of nuclear DNA amount in diploid bananas (Musa acuminata and M. balbisiana). Biol Plant 36: 351–357

Ducruet J-M, Peeva V, Havaux M (2007) Chlorophyll thermofluorescence and thermoluminescence as complementary tools for the study of temperature stress in plants. Photosynth Res 93: 159–171

Emms DM, Kelly S (2019) OrthoFinder: phylogenetic orthology inference for comparative genomics. Genome Biol 20: 238

Estravis-Barcala M, Heer K, Marchelli P, Ziegenhagen B, Arana MV, Bellora N (2021) Deciphering the transcriptomic regulation of heat stress responses in Nothofagus pumilio. PLOS ONE 16: e0246615

Ewels P, Magnusson M, Lundin S, Käller M (2016) MultiQC: summarize analysis results for multiple tools and samples in a single report. Bioinformatics 32: 3047–3048

Flynn JM, Hubley R, Goubert C, Rosen J, Clark AG, Feschotte C, Smit AF (2020) RepeatModeler2 for automated genomic discovery of transposable element families. Proc Natl Acad Sci U S A 117: 9451–9457

Geange SR, Arnold PA, Catling AA, Coast O, Cook AM, Gowland KM, Leigh A, Notarnicola RF, Posch BC, Venn SE, et al (2021) The thermal tolerance of photosynthetic tissues: a global systematic review and agenda for future research. New Phytol 229: 2497–2513

Gobiet A, Kotlarski S, Beniston M, Heinrich G, Rajczak J, Stoffel M (2014) 21st century climate change in the European Alps--a review. Sci Total Environ 493: 1138–1151

Guihur A, Rebeaud ME, Goloubinoff P (2022) How do plants feel the heat and survive? Trends Biochem Sci 47: 824–838

Gurevich A, Saveliev V, Vyahhi N, Tesler G (2013) QUAST: quality assessment tool for genome assemblies. Bioinformatics 29: 1072–1075

Hobbs JK, Jiao W, Easter AD, Parker EJ, Schipper LA, Arcus VL (2013) Change in heat capacity for enzyme catalysis determines temperature dependence of enzyme catalyzed rates. ACS Chem Biol 8: 2388–2393

Huang J, Zhao X, Bürger M, Wang Y, Chory J (2021) Two interacting ethylene response factors regulate heat stress response. Plant Cell 33: 338–357

Hugly S, Kunst L, Browse J, Somerville C (1989) Enhanced thermal tolerance of photosynthesis and altered chloroplast ultrastructure in a mutant of Arabidopsis deficient in lipid desaturation. Plant Physiol 90: 1134–1142

Ilík P, Kouril R, Kruk J, Myśliwa-Kurdziel B, Popelková H, Strzałka K, Naus J (2003) Origin of chlorophyll fluorescence in plants at 55-75 degrees C. Photochem Photobiol 77: 68–76

IPCC (2023) Climate Change 2021 – The Physical Science Basis: Working Group I Contribution to the Sixth Assessment Report of the Intergovernmental Panel on Climate Change. Cambridge University Press

Jiang Y, Feng X, Wang H, Chen Y, Sun Y (2021) Heat-induced down-regulation of photosystem II protects photosystem I in honeysuckle (Lonicera japonica). Journal of Plant Research 134: 1311–1321

Jones A, Torkel C, Stanley D, Nasim J, Borevitz J, Schwessinger B (2021) High-molecular weight DNA extraction, clean-up and size selection for long-read sequencing. PLoS One 16: e0253830

Kataoka R, Takahashi M, Suzuki N (2017) Coordination between bZIP28 and HSFA2 in the regulation of heat response signals in Arabidopsis. Plant Signal Behav 12: e1376159

Knight CA, Ackerly DD (2002) An ecological and evolutionary analysis of photosynthetic thermotolerance using the temperature-dependent increase in fluorescence. Oecologia 130: 505–514

Koren S, Walenz BP, Berlin K, Miller JR, Bergman NH, Phillippy AM (2017) Canu: scalable and accurate long-read assembly via adaptive-mer weighting and repeat separation. Genome Res 27: 722–736

Kramer MC, Janssen KA, Palos K, Nelson ADL, Vandivier LE, Garcia BA, Lyons E, Beilstein MA, Gregory BD (2020) N-methyladenosine and RNA secondary structure affect transcript stability and protein abundance during systemic salt stress in Arabidopsis. Plant Direct 4: e00239

Kriventseva EV, Kuznetsov D, Tegenfeldt F, Manni M, Dias R, Simão FA, Zdobnov EM (2019) OrthoDB v10: sampling the diversity of animal, plant, fungal, protist, bacterial and viral genomes for evolutionary and functional annotations of orthologs. Nucleic Acids Res 47: D807–D811

Kulkarni SR, Vaneechoutte D, Van de Velde J, Vandepoele K (2018) TF2Network: predicting transcription factor regulators and gene regulatory networks in Arabidopsis using publicly available binding site information. Nucleic Acids Res 46: e31

Langfelder P, Horvath S (2008) WGCNA: an R package for weighted correlation network analysis. BMC Bioinformatics 9: 559

Larkindale J, Huang B (2004) Thermotolerance and antioxidant systems in Agrostis stolonifera: involvement of salicylic acid, abscisic acid, calcium, hydrogen peroxide, and ethylene. J Plant Physiol 161: 405–413

Liao Y, Smyth GK, Shi W (2013) The Subread aligner: fast, accurate and scalable read mapping by seed-and-vote. Nucleic Acids Res 41: e108

Li H (2018) Minimap2: pairwise alignment for nucleotide sequences. Bioinformatics 34: 3094–3100

Li L, Duncan O, Ganguly DR, Lee CP, Crisp PA, Wijerathna-Yapa A, Salih K, Trösch J, Pogson BJ, Millar AH (2022) Enzymes degraded under high light maintain proteostasis by transcriptional regulation in. Proc Natl Acad Sci U S A 119: e2121362119

Li N, Euring D, Cha JY, Lin Z, Lu M, Huang L-J, Kim WY (2020a) Plant Hormone-Mediated Regulation of Heat Tolerance in Response to Global Climate Change. Front Plant Sci 11: 627969

Lippmann R, Babben S, Menger A, Delker C, Quint M (2019) Development of Wild and Cultivated Plants under Global Warming Conditions. Curr Biol 29: R1326–R1338

Liu J, Sun N, Liu M, Liu J, Du B, Wang X, Qi X (2013) An autoregulatory loop controlling Arabidopsis HsfA2 expression: role of heat shock-induced alternative splicing. Plant Physiol 162: 512–521

Liu R, Holik AZ, Su S, Jansz N, Chen K, Leong HS, Blewitt ME, Asselin-Labat M-L, Smyth GK, Ritchie ME (2015) Why weight? Modelling sample and observational level variability improves power in RNA-seq analyses. Nucleic Acids Res 43: e97

Li Z, Tang J, Srivastava R, Bassham DC, Howell SH (2020b) The Transcription Factor bZIP60 Links the Unfolded Protein Response to the Heat Stress Response in Maize. Plant Cell 32: 3559–3575

Marçais G, Kingsford C (2011) A fast, lock-free approach for efficient parallel counting of occurrences of k-mers. Bioinformatics 27: 764–770

Mittler R, Finka A, Goloubinoff P (2012) How do plants feel the heat? Trends Biochem Sci 37: 118–125

Nicotra AB, Segal DL, Hoyle GL, Schrey AW, Verhoeven KJF, Richards CL (2015) Adaptive plasticity and epigenetic variation in response to warming in an Alpine plant. Ecol Evol 5: 634–647

Niu Y, Xiang Y (2018) An Overview of Biomembrane Functions in Plant Responses to High-Temperature Stress. Front Plant Sci 9: 915

Notarnicola RF, Nicotra AB, Kruuk LEB, Arnold PA (2021) Tolerance of Warmer Temperatures Does Not Confer Resilience to Heatwaves in an Alpine Herb. Front Ecol Evol 9: 615119

Notarnicola RF, Nicotra AB, Kruuk LEB, Arnold PA (2023) Effects of warming temperatures on germination responses and trade-offs between seed traits in an alpine plant. Journal of Ecology 111: 62–76

Nurk S, Walenz BP, Rhie A, Vollger MR, Logsdon GA, Grothe R, Miga KH, Eichler EE, Phillippy AM, Koren S (2020) HiCanu: accurate assembly of segmental duplications, satellites, and allelic variants from high-fidelity long reads. Genome Res 30: 1291–1305

Oomen RA, Hutchings JA (2022) Genomic reaction norms inform predictions of plastic and adaptive responses to climate change. J Anim Ecol 91: 1073–1087

Oomen RA, Hutchings JA (2017) Transcriptomic responses to environmental change in fishes: Insights from RNA sequencing. FACETS. doi: 10.1139/facets-2017-0015

Pertea G, Pertea M (2020) GFF Utilities: GffRead and GffCompare. F1000Res. doi: 10.12688/f1000research.23297.2

de Pinto MC, Locato V, Paradiso A, De Gara L (2015) Role of redox homeostasis in thermo-tolerance under a climate change scenario. Ann Bot 116: 487–496

Poór P, Nawaz K, Gupta R, Ashfaque F, Khan MIR (2022) Ethylene involvement in the regulation of heat stress tolerance in plants. Plant Cell Rep 41: 675–698

Posch BC, Amoanimaa-Dede H, Aparecido LMT, Atkin OK, Bison NN, Blonder BW, Coast O, Doughty CE, Guo JS, van Haren J, et al (2026) High-temperature acclimation of photosystem II in land plants. New Phytol 249: 1108–1123

Quinlan AR, Hall IM (2010) BEDTools: a flexible suite of utilities for comparing genomic features. Bioinformatics 26: 841–842

Ranallo-Benavidez TR, Jaron KS, Schatz MC (2020) GenomeScope 2.0 and Smudgeplot for reference-free profiling of polyploid genomes. Nat Commun 11: 1432

Rashid FAA, Crisp PA, Zhang Y, Berkowitz O, Pogson BJ, Day DA, Masle J, Dewar RC, Whelan J, Atkin OK, et al (2020) Molecular and physiological responses during thermal acclimation of leaf photosynthesis and respiration in rice. Plant Cell Environ 43: 594–610

Reddy ASN, Marquez Y, Kalyna M, Barta A (2013) Complexity of the alternative splicing landscape in plants. Plant Cell 25: 3657–3683

Rey O, Danchin E, Mirouze M, Loot C, Blanchet S (2016) Adaptation to Global Change: A Transposable Element-Epigenetics Perspective. Trends Ecol Evol 31: 514–526

Richter K, Haslbeck M, Buchner J (2010) The heat shock response: life on the verge of death. Mol Cell 40: 253–266

Ritchie ME, Phipson B, Wu D, Hu Y, Law CW, Shi W, Smyth GK (2015) limma powers differential expression analyses for RNA-sequencing and microarray studies. Nucleic Acids Res 43: e47

Robinson MD, Oshlack A (2010) A scaling normalization method for differential expression analysis of RNA-seq data. Genome Biol 11: R25

Scafaro AP, Bautsoens N, den Boer B, Van Rie J, Gallé A (2019) A Conserved Sequence from Heat-Adapted Species Improves Rubisco Activase Thermostability in Wheat. Plant Physiol 181: 43–54

Schreiber U, Berry JA (1977) Heat-induced changes of chlorophyll fluorescence in intact leaves correlated with damage of the photosynthetic apparatus. Planta 136: 233–238

Seppey M, Manni M, Zdobnov EM (2019) BUSCO: Assessing Genome Assembly and Annotation Completeness. Methods Mol Biol 1962: 227–245

Shen W, Sipos B, Zhao L (2024) SeqKit2: A Swiss army knife for sequence and alignment processing. Imeta 3: e191

Soneson C, Love MI, Robinson MD (2015) Differential analyses for RNA-seq: transcript-level estimates improve gene-level inferences. F1000Res 4: 1521

Sritharan MS, Hemmings FA, Moles AT (2021) Few changes in native Australian alpine plant morphology, despite substantial local climate change. Ecol Evol 11: 4854–4865

Stanke M, Schöffmann O, Morgenstern B, Waack S (2006) Gene prediction in eukaryotes with a generalized hidden Markov model that uses hints from external sources. BMC Bioinformatics 7: 62

Sun J-L, Li J-Y, Wang M-J, Song Z-T, Liu J-X (2021) Protein Quality Control in Plant Organelles: Current Progress and Future Perspectives. Mol Plant 14: 95–114

Vandersteen Tymchuk W, O’Reilly P, Bittman J, Macdonald D, Schulte P (2010) Conservation genomics of Atlantic salmon: variation in gene expression between and within regions of the Bay of Fundy. Mol Ecol 19: 1842–1859

Wang L, Ma K-B, Lu Z-G, Ren S-X, Jiang H-R, Cui J-W, Chen G, Teng N-J, Lam H-M, Jin B (2020) Differential physiological, transcriptomic and metabolomic responses of Arabidopsis leaves under prolonged warming and heat shock. BMC Plant Biol 20: 86

Wang W, Teng F, Lin Y, Ji D, Xu Y, Chen C, Xie C (2018) Transcriptomic study to understand thermal adaptation in a high temperature-tolerant strain of Pyropia haitanensis. PLoS One 13: e0195842

Way DA, Yamori W (2014) Thermal acclimation of photosynthesis: on the importance of adjusting our definitions and accounting for thermal acclimation of respiration. Photosynth Res 119: 89–100

Wu Y-S, Yang C-Y (2019) Ethylene-mediated signaling confers thermotolerance and regulates transcript levels of heat shock factors in rice seedlings under heat stress. Bot Stud 60: 23

Yamori W, Hikosaka K, Way DA (2014) Temperature response of photosynthesis in C3, C4, and CAM plants: temperature acclimation and temperature adaptation. Photosynth Res 119: 101–117

Zhang B, Horvath S (2005) A general framework for weighted gene co-expression network analysis. Stat Appl Genet Mol Biol 4: Article17

Zivcak M, Brestic M, Kalaji HM, Govindjee (2014) Photosynthetic responses of sun- and shade-grown barley leaves to high light: is the lower PSII connectivity in shade leaves associated with protection against excess of light? Photosynth Res 119: 339–354

