## Supplementary Tables S1-S3; Supplementary Figures S1-S3 for "Molecular signatures of heat tolerance in an Australian alpine plant during moderate warming"

Plant Physiology

**List of Supplementary Tables**

- Supplementary Data 1: mRNA sequencing statistics
- Supplementary Data 2: Orthogroups identified between Wahlenbergia ceracea and Arabidopsis thaliana by OrthoFinder
- Supplementary Data 3: Differentially expressed genes in response to warming across all lines
- Supplementary Data 4: Differentially expressed genes in response to warming in tolerant lines
- Supplementary Data 5: Differentially expressed genes in response to warming in sensitive lines
- Supplementary Data 6: Arabidopsis orthologs of warming-induced genes in *Wahlenbergia ceracea*
- Supplementary Data 7: Co-expressed gene modules identified by WGCNA
- Supplementary Data 8: Gene membership for significant module-trait correlations identified from WGCNA
- Supplementary Data 9: Constructing gene regulatory networks and identifying cognate transcription factors using TF2Network

**Table S1.** Pollen doner and recipient history to generate *W. ceracea* lines used in this study.

| $F_1$ generation | | | $F_2$ generation | | | $F_3$ generation | | |
|---|---|---|---|---|---|---|---|---|
| Recipient ID | Donor ID | ID | Recipient ID | Donor ID | ID | Recipient ID | Donor ID | ID |
| $F_0$.1636 | $F_0$.1654 | $F_1$.128 | $F_1$.128 | $F_1$.111 | $F_2$.214 | $F_2$.214 | $F_2$.223 | $F_3$.485 |
| $F_0$.1504 | $F_0$.1501 | $F_1$.120 | $F_1$.120 | $F_1$.111 | $F_2$.216 | $F_2$.216 | $F_2$.225 | $F_3$.491 |
| $F_0$.1653 | $F_0$.1523 | $F_1$.144 | $F_1$.144 | $F_1$.148 | $F_2$.244 | $F_2$.244 | $F_2$.225 | $F_3$.495 |
| $F_0$.1623 | $F_0$.1630 | $F_1$.141 | $F_1$.141 | $F_1$.142 | $F_2$.240 | $F_2$.240 | $F_2$.217 | $F_3$.479 |
| $F_0$.1620 | $F_0$.1614 | $F_1$.103 | $F_1$.103 | $F_1$.111 | $F_2$.261 | $F_2$.261 | $F_2$.209 | $F_3$.467 |
| $F_0$.1523 | $F_0$.1511 | $F_1$.147 | $F_1$.147 | $F_1$.127 | $F_2$.230 | $F_2$.230 | $F_2$.217 | $F_3$.483 |
| $F_0$.1651 | $F_0$.1509 | $F_1$.148 | $F_1$.148 | $F_1$.127 | $F_2$.255 | $F_2$.255 | $F_2$.201 | $F_3$.457 |
| $F_0$.1643 | $F_0$.1639 | $F_1$.133 | $F_1$.144 | $F_1$.101 | $F_2$.249 | $F_2$.249 | $F_2$.217 | $F_3$.439 |
| $F_0$.1618 | $F_0$.1616 | $F_1$.126 | $F_1$.133 | $F_1$.148 | $F_2$.242 | $F_2$.242 | $F_2$.213 | $F_3$.475 |
| $F_0$.1537 | $F_0$.1603 | $F_1$.101 | $F_1$.126 | $F_1$.142 | $F_2$.257 | $F_2$.257 | $F_2$.223 | $F_3$.487 |
| $F_0$.1646 | $F_0$.1641 | $F_1$.149 | $F_1$.101 | $F_1$.123 | $F_2$.224 | $F_2$.224 | $F_2$.213 | $F_3$.477 |
| $F_0$.1509 | $F_0$.1514 | $F_1$.117 | $F_1$.149 | $F_1$.153 | $F_2$.264 | $F_2$.264 | $F_2$.213 | $F_3$.437 |
| $F_0$.1629 | $F_0$.1526 | $F_1$.119 | $F_1$.117 | $F_1$.123 | $F_2$.223 | | | |
| $F_0$.1543 | $F_0$.1534 | $F_1$.130 | $F_1$.119 | $F_1$.126 | $F_2$.225 | | | |
| $F_0$.1525 | $F_0$.1524 | $F_1$.115 | $F_1$.130 | $F_1$.118 | $F_2$.217 | | | |
| $F_0$.1533 | $F_0$.1612 | $F_1$.114 | $F_1$.115 | $F_1$.109 | $F_2$.209 | | | |
| $F_0$.1511 | $F_0$.1513 | $F_1$.107 | $F_1$.114 | $F_1$.101 | $F_2$.201 | | | |
| $F_0$.1602 | $F_0$.1536 | $F_1$.111 | $F_1$.107 | $F_1$.111 | $F_2$.213 | | | |
| $F_0$.1643 | $F_0$.1655 | $F_1$.142 | | | | | | |
| $F_0$.1648 | $F_0$.1639 | $F_1$.127 | | | | | | |
| $F_0$.1508 | $F_0$.1502 | $F_1$.123 | | | | | | |
| $F_0$.1507 | $F_0$.1501 | $F_1$.153 | | | | | | |
| $F_0$.1605 | $F_0$.1610 | $F_1$.118 | | | | | | |
| $F_0$.1613 | $F_0$.1606 | $F_1$.109 | | | | | | |

**Table S2.** Summary of annotated repeats identified in our genome assembly for *W. ceracea*.

| Class | Subclass/Family | Number of elements | Length occupied (bp) | Percentage of Assembly |
|---|---|---|---|---|
| **Retroelements** | | **666,153** | **563,084,583** | **41.22** |
| | **LINEs** | **16,271** | **6,314,206** | **0.46** |
| | RTE/Bov-B | 2,109 | 356,251 | 0.03 |
| | L1/CIN4 | 14,162 | 5,957,955 | 0.44 |
| | **LTR elements** | **649,882** | **556,770,377** | **40.76** |
| | Ty1/Copia | 217,468 | 212,171,636 | 15.53 |
| | Gypsy/DIRS1 | 240,192 | 269,849,111 | 19.76 |
| **DNA Transposons** | | **74,845** | **56,515,625** | **4.14** |
| | hobo-Activator | 2,580 | 1,773,619 | 0.13 |
| | Tc1-IS630-Pogo | 1,988 | 708,358 | 0.05 |
| | MULE-MuDR | 40,029 | 23,969,531 | 1.75 |
| | Tourist/Harbinger | 18,692 | 12,754,736 | 0.93 |
| | **Rolling-circles** | **9,144** | **4,070,659** | **0.30** |
| | **Unclassified** | **758,988** | **367,422,193** | **26.90** |
| **Total Interspersed Repeats** | | | **987,022,401** | **72.26** |
| | Small RNA | 6,827 | 14,259,421 | 1.04 |
| | Simple repeats | 328,912 | 26,335,934 | 1.93 |
| | Low complexity | 38,636 | 1,855,441 | 0.14 |
| **Total soft-masked** | | | **1,033,543,856** | **75.67** |

**Table S3.** Number of orthogroups between species. *W. cer* = *Wahlenbergia ceracea*; *M. esc* = *Manihot esculenta*; *G. max* = *Glycine max*; *V. vin* = *Vitis vinifera*; *S. lyc* = *Solanum lycopersicum*; *E. gra* = *Eucalyptus grandis*; *A. tha* = *Arabidopsis thaliana*; *O. sat* = *Oryza sativa*; *Z. mays* = *Zea mays*; *M. pol* = *Marchantia polymorpha*; *S. moe* = *Selaginella moellendorffii*; *P. pat* = *Physcomitrium patens*; *C. rei* = *Chlamydomonas reinhardtii*.

| | *A. tha* | *C. reii* | *E. gra* | *G. max* | *M. esc* | *M. pol* | *O. sat* | *P. pat* | *S. moe* | *S. lyc* | *V. vin* | *Z. mays* | *W. cer* |
|---|---|---|---|---|---|---|---|---|---|---|---|---|---|
| *W. cer* | 8854 | 4345 | 9060 | 9275 | 9357 | 6400 | 8474 | 6265 | 6341 | 9165 | 9239 | 8332 | 11481 |
| *M. esc* | 10089 | 4710 | 10314 | 10711 | 12184 | 6997 | 9452 | 6837 | 6864 | 10234 | 10490 | 9298 | 9357 |
| *G. max* | 10059 | 4716 | 10308 | 13760 | 10711 | 6998 | 9503 | 6872 | 6900 | 10201 | 10451 | 9371 | 9275 |
| *V. vin* | 9844 | 4685 | 10178 | 10451 | 10490 | 6955 | 9431 | 6802 | 6867 | 10078 | 12044 | 9272 | 9239 |
| *S. lyc* | 9724 | 4615 | 9864 | 10201 | 10234 | 6844 | 9238 | 6713 | 6758 | 11925 | 10078 | 9061 | 9165 |
| *E. gra* | 9773 | 4641 | 12155 | 10308 | 10314 | 6876 | 9324 | 6703 | 6795 | 9864 | 10178 | 9210 | 9060 |
| *A. tha* | 12082 | 4684 | 9773 | 10059 | 10089 | 6910 | 9158 | 6770 | 6795 | 9724 | 9844 | 9043 | 8854 |
| *O. sat* | 9158 | 4679 | 9324 | 9503 | 9452 | 6882 | 14499 | 6771 | 6794 | 9238 | 9431 | 11978 | 8474 |
| *Z. mays* | 9043 | 4649 | 9210 | 9371 | 9298 | 6805 | 11978 | 6680 | 6725 | 9061 | 9272 | 15328 | 8332 |
| *M. poly* | 6910 | 4983 | 6876 | 6998 | 6997 | 9531 | 6882 | 7552 | 6846 | 6844 | 6955 | 6805 | 6400 |
| *S. moe* | 6795 | 4726 | 6795 | 6900 | 6864 | 6846 | 6794 | 6770 | 10881 | 6758 | 6867 | 6725 | 6341 |
| *P. pat* | 6770 | 4978 | 6703 | 6872 | 6837 | 7552 | 6771 | 14690 | 6770 | 6713 | 6802 | 6680 | 6265 |
| *C. rei* | 4684 | 7171 | 4641 | 4716 | 4710 | 4983 | 4679 | 4978 | 4726 | 4615 | 4685 | 4649 | 4345 |

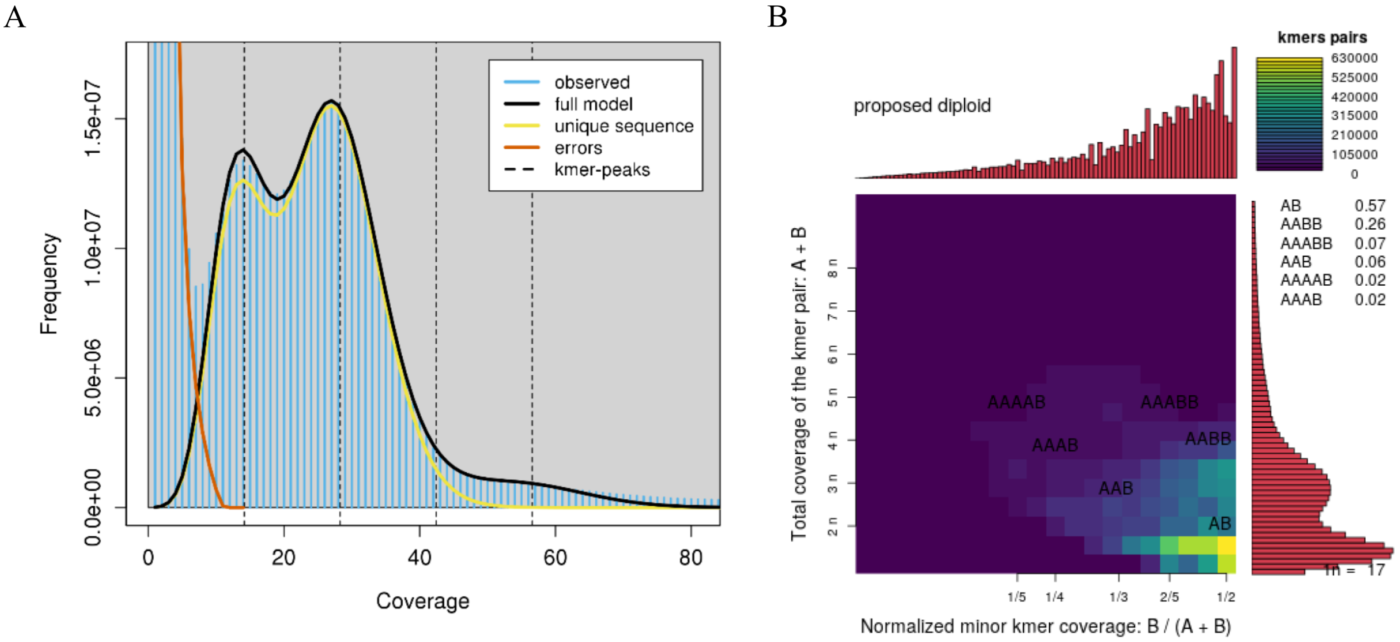


**Figure S1 Assessment of Ploidy**

(A) 31-mer spectra and fitted models from unassembled long-read data of *Wahlenbergia ceracea*. Modelled peaks corresponding to heterozygous and homozygous k-mer multiplicities (~14x, 28x, 42x, and 56x) suggest diploidy. Low-coverage 31-mers were classified as errors.

(B) Genotype distribution inferred from Smudgeplot analysis of 21-mers is consistent with diploidy.

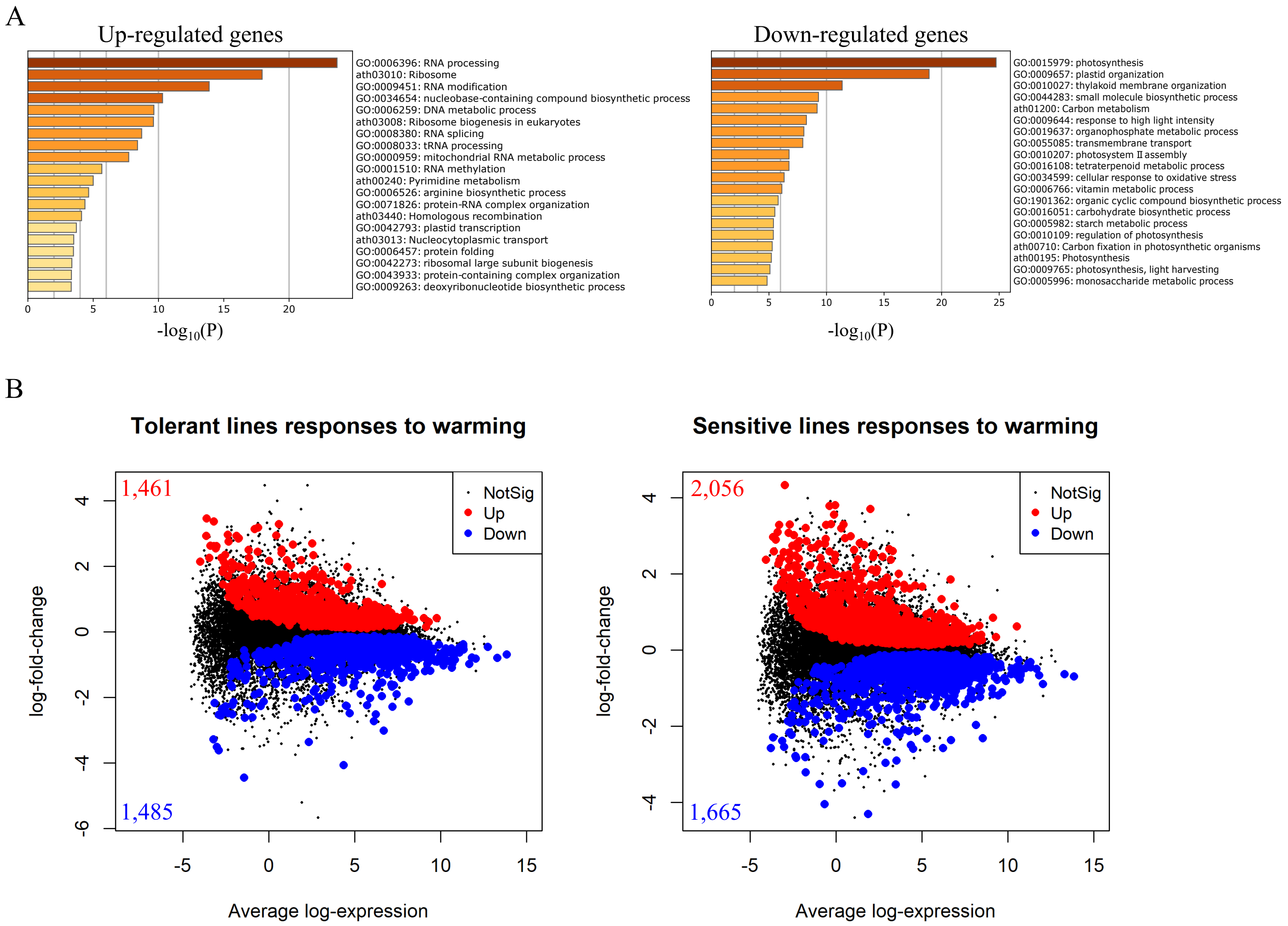


**Figure S2 Prolong moderate warming induces a robust transcriptional response in *W. ceracea***
(A) Bar charts representing the statistical significance of enriched functional terms for up- and down-regulated genes detected from Contrast *I*.
(B) Mean-abundance plots where each dot represents a gene, plotted by its average abundance against its $\log_2$ fold-change. Red and blue dots indicate differentially expressed genes in tolerant and sensitive *W. ceracea* lines (FDR < 0.05).

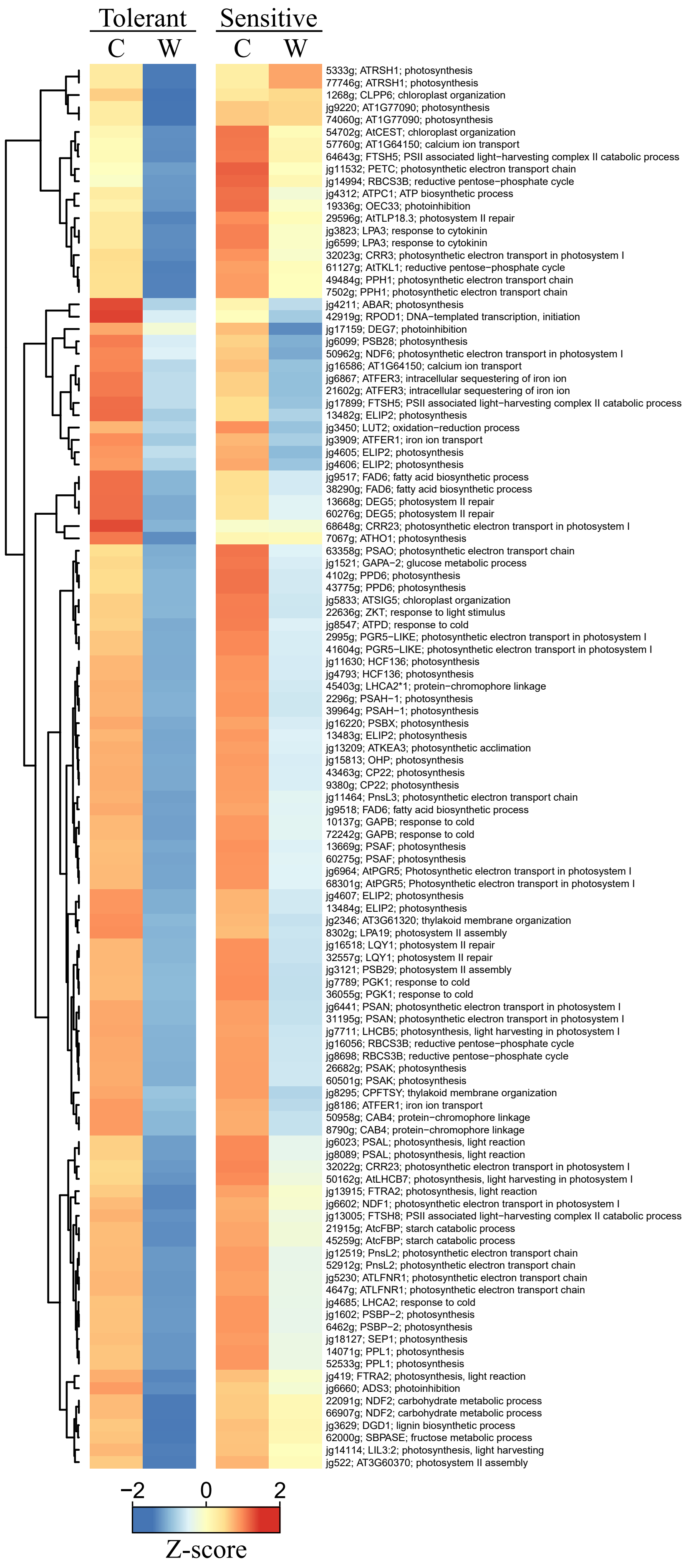


**Figure S3 Warming-induced differential expression of photosynthesis-associated genes**
Heatmap with one-dimensional hierarchical clustering of differentially expressed genes encoding photosynthesis-related proteins  Cell colour denotes relative expression values (row-wise Z-scores) under cool (C) and warm (W) conditions. Labels denote *W. ceracea* gene ID, Arabidopsis gene ID and name or function.
